# Gallionellaceae pangenomic analysis reveals insight into phylogeny, metabolic flexibility, and iron oxidation mechanisms

**DOI:** 10.1101/2023.01.26.525709

**Authors:** Rene L. Hoover, Jessica L. Keffer, Shawn W. Polson, Clara S. Chan

**Author notes:** Corresponding author: Clara S. Chan.

## Abstract

The iron-oxidizing Gallionellaceae drive a wide variety of biogeochemical cycles through their metabolisms and biominerals. To better understand the environmental impacts of Gallionellaceae, we need to improve our knowledge of their diversity and metabolisms, especially any novel iron oxidation mechanisms. Here, we used a pangenomic analysis of 103 genomes to resolve Gallionellaceae phylogeny and explore the range of genomic potential. Using a concatenated ribosomal protein tree and key gene patterns, we determined Gallionellaceae has four genera, divided into two groups–iron-oxidizing bacteria (FeOB) *Gallionella*, *Sideroxydans*, and *Ferriphaselus* with known iron oxidases (Cyc2, MtoA) and nitrite-oxidizing bacteria (NOB) *Candidatus* Nitrotoga with nitrite oxidase (Nxr). The FeOB and NOB have similar electron transport chains, including genes for reverse electron transport and carbon fixation. Auxiliary energy metabolisms including S oxidation, denitrification, and organotrophy were scattered throughout the Gallionellaceae FeOB. Within FeOB, we found genes that may represent adaptations for iron oxidation, including a variety of extracellular electron uptake (EEU) mechanisms. FeOB genomes encoded more predicted *c*-type cytochromes overall, notably more multiheme *c*-type cytochromes (MHCs) with >10 CXXCH motifs. These include homologs of several predicted outer membrane porin-MHC complexes, including MtoAB and Uet. MHCs are known to efficiently conduct electrons across longer distances and function across a wide range of redox potentials that overlap with mineral redox potentials, which can help expand the range of usable iron substrates. Overall, the results of pangenome analyses suggest that the Gallionellaceae genera *Gallionella*, *Sideroxydans*, and *Ferriphaselus* are primarily iron oxidizers, capable of oxidizing dissolved Fe^2+^ as well as a range of solid iron or other mineral substrates.

**Importance:** Neutrophilic iron-oxidizing bacteria (FeOB) produce copious iron (oxyhydr)oxides that can profoundly influence biogeochemical cycles, notably the fate of carbon and many metals. To fully understand environmental microbial iron oxidation, we need a thorough accounting of iron oxidation mechanisms. In this study we show the Gallionellaceae FeOB have both known iron oxidases as well as uncharacterized multiheme cytochromes (MHCs). MHCs are predicted to transfer electrons from extracellular substrates and likely confer metabolic capabilities that help Gallionellaceae occupy a range of different iron- and mineral-rich niches. Gallionellaceae appear to specialize in iron oxidation, so it makes sense that they would have multiple mechanisms to oxidize various forms of iron, given the many iron minerals on Earth, as well as the physiological and kinetic challenges faced by FeOB. The multiple iron/mineral oxidation mechanisms may help drive the widespread ecological success of Gallionellaceae.

## Introduction

*Gallionella* are one of the oldest known and most well studied iron-oxidizing bacteria (FeOB), yet we are still learning how they oxidize iron and adapt to iron-rich niches. *Gallionella* is the type genus of the family Gallionellaceae, which also includes *Sideroxydans*, *Ferriphaselus*, and *Ferrigenium*. These Gallionellaceae FeOB are found in a wide range of environments, including freshwater creeks, sediment, root rhizospheres, peat, permafrost, deep subsurface aquifers, and municipal waterworks (1–18). FeOB potentially drive the fate of many metals and nutrients via both metabolic reactions and forming iron oxyhydroxides that adsorb and react with many solutes (19). To better understand the biogeochemical effects of Gallionellaceae, we need to improve our knowledge of their phylogeny and metabolic mechanisms, especially for iron oxidation. Recently, the rapid increase in metagenomes from iron-rich environments has significantly expanded the number of available Gallionellaceae genomes, which makes it possible to investigate diversity and mechanisms using genomic analyses of both cultured and uncultured Gallionellaceae.

The Gallionellaceae are named after *Gallionella ferruginea*, first described by Ehrenberg in 1838 (20), and recognizable by its distinctive, twisted iron oxyhydroxide stalk (21). While the type strain, *G. ferruginea* Johan (22) no longer exists, there are seven iron-oxidizing Gallionellaceae isolates (7, 11, 23–26). Some isolates, such as *Ferriphaselus* spp., appear to be obligate iron oxidizers, while others also grow on non-iron substrates. In addition to iron, *S. lithotrophicus* ES-1 grows by thiosulfate oxidation (24, 27) while *Sideroxydans* sp. CL21 shows mixotrophic growth with either lactate or hydrogen (28). Some *Ferrigenium* can also reduce nitrate (29, 30). It is unknown how common it is for Gallionellaceae to use electron donors/acceptors besides Fe(II)/O_2_, though these alternate metabolisms may help their success across different environments and fluctuating conditions typical of many oxic-anoxic interfaces. Even so, since all seven Gallionellaceae isolates are neutrophilic aerobic chemolithoautotrophic iron oxidizers, this could be the dominant metabolic niche of Gallionellaceae.

In Gallionellaceae and other neutrophilic chemolithotrophic FeOB, there are two known iron oxidases: Cyc2, a fused monoheme cytochrome-porin and MtoAB, a decaheme porin-cytochrome complex (31–33). The *mtoA* (<u>m</u>etal <u>o</u>xidation) gene was first identified and characterized in FeOB *S. lithotrophicus* ES-1 (31). The *mtoA* gene is a homolog of both *pioA* (<u>p</u>hototrophic iron <u>o</u>xidation), which encodes the PioA iron oxidase in the photoferrotroph *Rhodopseudomonas palustris* TIE-1 (34, 35), and *mtrA* (<u>m</u>etal reduction), which encodes the MtrA iron reductase in iron-reducing bacteria (FeRB) *Shewanella* (36). The *cyc2* gene is more common than *mtoAB* and is found in nearly all well-characterized neutrophilic FeOB like the Gallionellaceae (32) and Zetaproteobacteria (33), making it a suitable genetic marker for many FeOB. Cyc2 has been demonstrated to oxidize aqueous Fe^2+^ (32), while Mto gene/protein expression has been associated with the oxidation of solid iron minerals (37). However, Cyc2 and MtoA may not be the only mechanisms for neutrophilic iron oxidation. There are a number of additional uncharacterized cytochromes and electron transport genes (27, 38) within Gallionellaceae genomes such as isolate *S. lithotrophicus* ES-1 (27, 38), suggesting the existence of novel iron oxidation genes and mechanisms within the family.

The Gallionellaceae also includes a recently identified genus, *Candidatus* Nitrotoga, which are chemolithotrophic nitrite-oxidizing bacteria (NOB). Like the iron-oxidizing Gallionellaceae, they are widespread in freshwater and engineered environments, including permafrost (39), coastal sediments (40), freshwater (41), freshwater sediments (42), and the activated sludge of wastewater treatment facilities (43, 44). There are only two isolates, *Ca.* Nitrotoga fabula (43) and *Ca.* Nitrotoga sp. AM1P (45), along with four genomes from enrichment cultures (42). *Ca.* Nitrotoga are adapted to niches with low nitrite, and oxidize it using a distinct high-affinity Nxr nitrite oxidoreductase (39, 42, 46). Extensive iron uptake mechanisms have been detected in *Ca.* Nitrotoga genomes, indicating the importance of iron for growth, likely due to the FeS cluster of Nxr (42). However, neither the isolates nor enrichments are known to oxidize Fe(II). If *Ca.* Nitrotoga lack the capacity to oxidize iron, then we can investigate the iron-oxidizing mechanisms and adaptations of Gallionellaceae through a comparative genomic analysis of iron-versus nitrite-oxidizing members.

Toward this goal, we took advantage of the growing number of environmental metagenomes and collected 103 high quality Gallionellaceae genomes and metagenome assembles genomes (MAGs). We used those sequences to resolve the Gallionellaceae phylogeny and delineate groups of iron and nitrite oxidizers. We searched for known and novel iron oxidation genes, other energy and nutrient metabolisms, and genes found exclusively in FeOB that may represent adaptations for an iron-oxidizing lifestyle. This work increases our understanding of Gallionellaceae family phylogeny and the metabolic traits of its genera. It also highlights some of the key multiheme cytochromes in Gallionellaceae FeOB, which may facilitate extracellular electron uptake (EEU) and the oxidation of different iron substrates.

## Results

### Phylogeny

We collected 103 high quality Gallionellaceae isolate genomes and metagenome assembled genomes (MAGs) from various databases and collections (Table S1). Many of these MAGs were only classified at the family level. To resolve the phylogeny, verify existing classifications, and classify the unknown Gallionellaceae, we constructed a concatenated protein tree (Figure 1) from 13 ribosomal protein sequences. Organisms in the tree formed distinct, well-supported clades that corresponded to the major genera: *Gallionella*, *Sideroxydans*, *Ferriphaselus*, and *Ca.* Nitrotoga (Figure 1). Most of the MAGs previously classified as Gallionellaceae and Gallionellales were found to be either *Gallionella* or *Sideroxydans*, with the exception of one that clustered with the *Ca*. Nitrotoga (*Ca*. Nitrotoga SL_21). Although some genomes formed sub-clades, many were organized along a continuum. Near the base of the *Gallionella* are *Ferrigenium kumadai* An22 and the nitrate-reducing iron-oxidizing bacteria (NRFeOB) of the Straub (KS) and Bremen Pond (BP) enrichments (Figure 1). There is not a clear boundary between the *Gallionella* and the relatively new *Ferrigenium* genus, so we included the *Ferrigenium* and NRFeOB with the *Gallionella* grouping for our analyses. We also constructed a 16S rRNA gene tree containing 24 sequences in our dataset along with 941 high-quality, full-length Gallionellaceae sequences from the SILVA database (Fig. S1), but bootstrap support was weaker and clades were less clearly resolved. Therefore, concatenated ribosomal proteins are a more reliable determinant of Gallionellaceae phylogeny than 16S rRNA genes.

**FIGURE 1.**
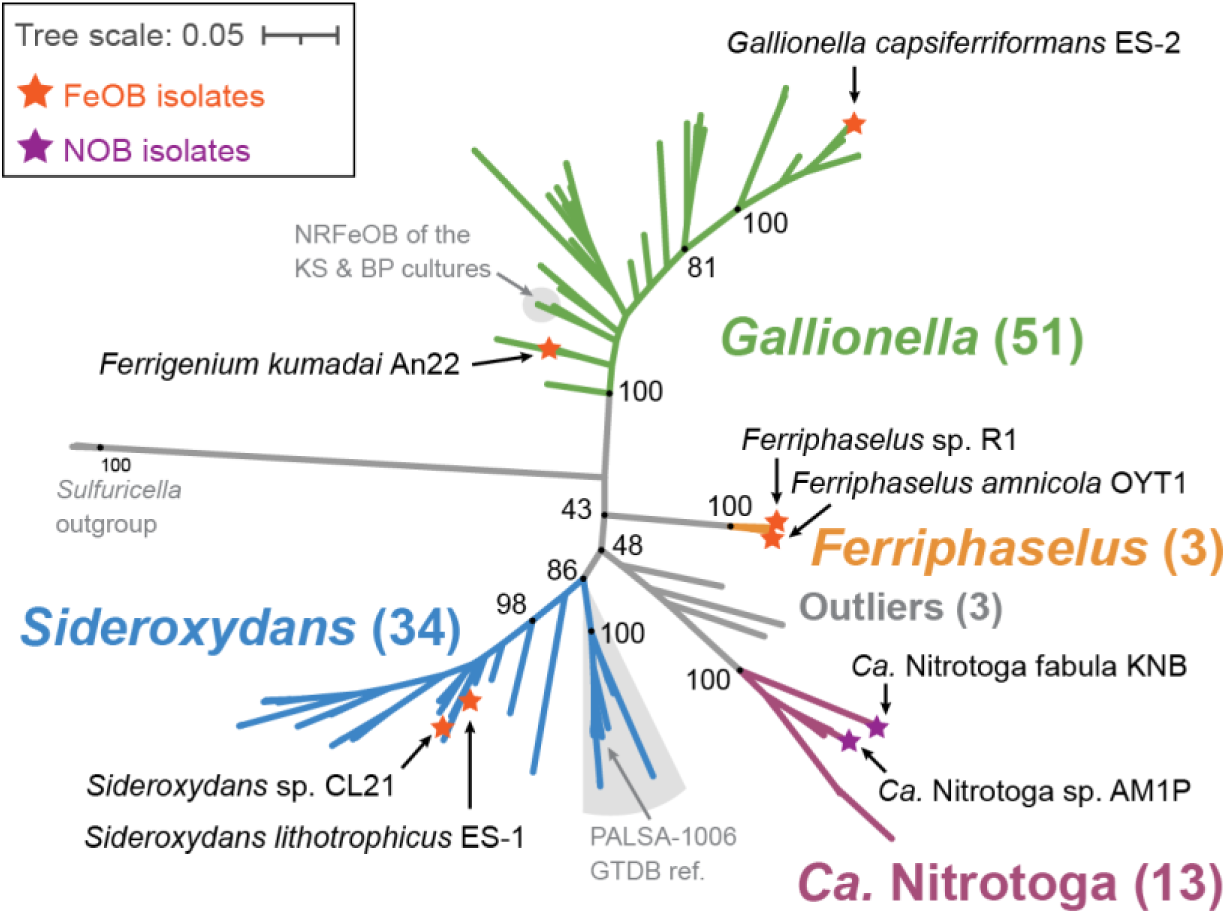
Concatenated ribosomal protein maximum likelihood tree of the Gallionellaceae family showing the four distinct genera: *Gallionella, Sideroxydans, Ferriphaselus*, and *Ca.* Nitrotoga. Isolates are labeled and annotated with stars. Support values from 1000 bootstraps shown for major branching nodes (black dots). Detailed tree shown in Fig. 2.

We assessed whether there was a relationship between phylogeny and environment. Each genome and MAG was classified with the GOLD classification schema (47) based on pre-existing GOLD classifications, available metadata and publications (Fig. 2, Table S2). The majority of aquatic genomes were from freshwater and groundwater environments while terrestrial genomes were mostly found in soil, peat, and rhizosphere environments. However, some genomes were sequenced from more extreme environments such as thermal hot springs (ENVO:00002181) and acid mine drainage (ENVO:00001997) (Table S2). Gallionellaceae are widespread and can inhabit many different environments, but there was no clear pattern between GOLD Ecosystem Type and broad phylogenetic groupings (Fig. 2). Different Gallionellaceae appear to co-exist in some environments, suggesting niches not captured in the ecosystem classification are controlling Gallionellaceae diversity and environmental distribution.

**FIGURE 2.**
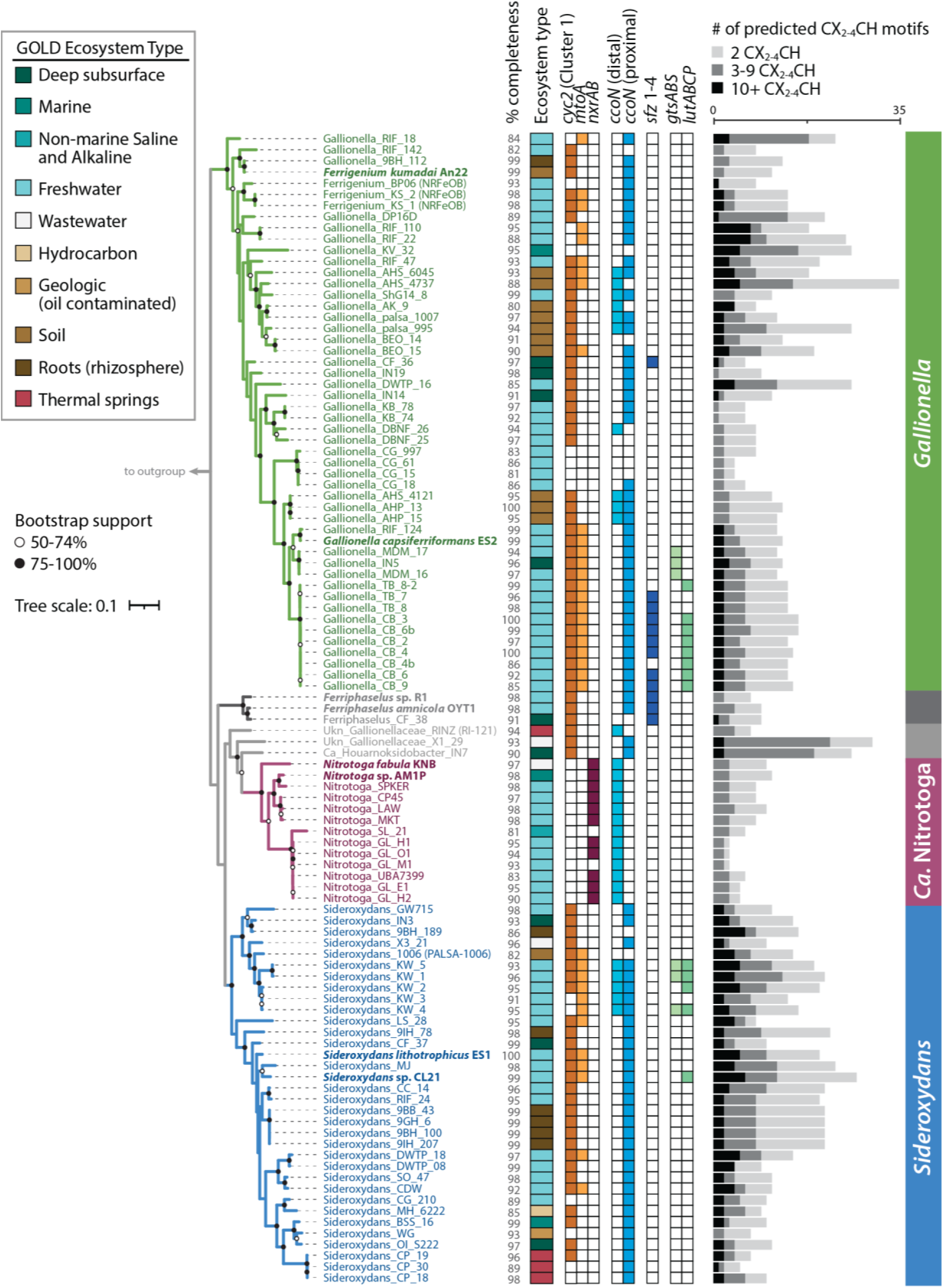
Maximum likelihood tree of concatenated ribosomal proteins from the Gallionellaceae annotated with source ecosystem and genes for iron oxidases (*cyc2, mtoA*), nitrite oxidase (*nxrAB*), terminal oxidase (*ccoN*), stalk formation (*sfz*), and organic utilization (*gtsABS*, *lutABCP*). The bar graph to the right shows the number multiheme cytochromes CXXCH, CX_3_CH, and CX_4_CH heme-binding motifs. Phylogeny does not correlate to environments, and key genes, including those for multiheme cytochromes, show distinct distributions between iron and nitrite oxidizer clades. Isolates are shown in bold. % completeness = genome completeness calculated with CheckM. Outgroup omitted for space.

### Metabolic potential and diversity

The Gallionellaceae family has few isolates, so to uncover the shared metabolic traits of its FeOB members, we compared and contrasted *Gallionella*, *Sideroxydans*, and *Ferriphaselus* genomes to those of the nitrite-oxidizing *Ca.* Nitrotoga. We identified key genes within the pangenome for iron oxidation (including predicted *c*-type cytochromes), carbon fixation, and respiration using a combination of DRAM (48), FeGenie (49), MagicLamp (50), a heme motif counter script (51), and BLAST (52, 53). To further uncover genes and pathways specifically enriched in the iron oxidizers, we used Anvi’o (54–56) to analyze a filtered dataset of only *Gallionella*, *Sideroxydans*, and *Ca*. Nitrotoga genomes that were >97% complete. This approach enabled us to create a comprehensive picture of Gallionellaceae metabolic diversity and pinpoint promising gene clusters that may be adaptations for an iron-oxidizing lifestyle.

### Primary energy metabolisms — iron and nitrite oxidation

Known metabolisms for the few Gallionellaceae isolates suggest *Ca*. Nitrotoga are nitrite oxidizers, while *Sideroxydans*, *Ferriphaselus*, and *Gallionella* are iron oxidizers. We examined the pangenome for the presence of *cyc2* and *mtoA* iron oxidase genes and *nxrAB* nitrite oxidase genes to determine if that pattern also holds throughout the uncultured Gallionellaceae. As with the isolates, there is a clear delineation between organisms with marker genes for iron versus nitrite oxidation, which corresponds to the phylogenetic groups (Fig. 2, Fig. 3).

**FIGURE 3.**
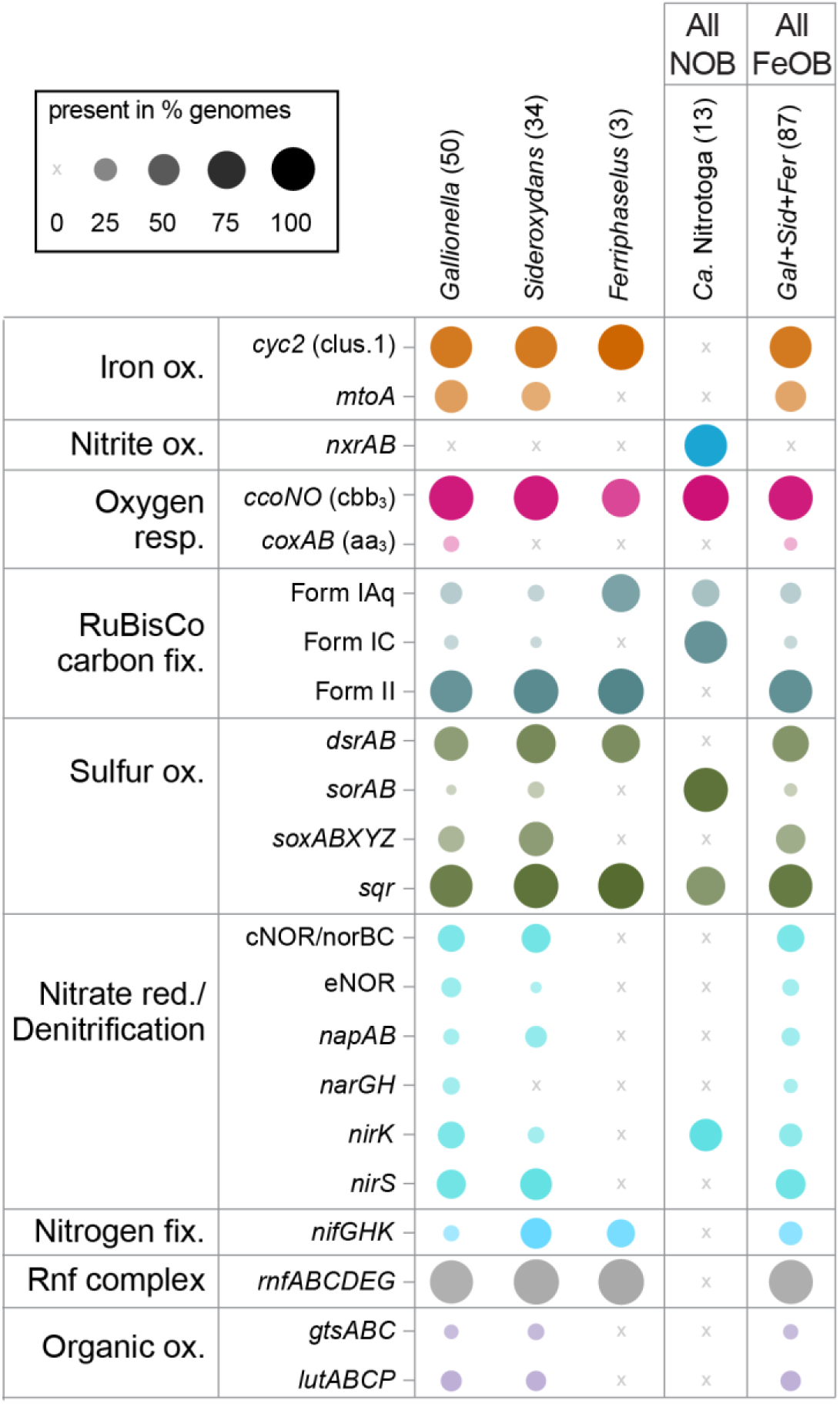
Plot showing the percent of genomes in each genus/group with genes for key metabolic pathways. The plot indicates the Gallionellaceae are aerobic lithoautotrophs with two main energy metabolisms, iron or nitrite oxidation. Some members also have metabolic potential for S oxidation and/or denitrification. Numbers in parentheses indicate the total number of genomes in each group. Color is used to distinguish groups, while dot size and opacity indicate % presence in the genome groups.

The *cyc2* gene is widespread among clades of iron oxidizers, with at least one copy detected in 83% of the FeOB genomes (Fig. 3, Table S3). The *mtoA* gene is found in 41% of the FeOB genomes, and 37% of genomes have both *mtoA* and *cyc2*. In total, 89% have at least one iron oxidase gene, either *cyc2* or *mtoA* (Table S3). Since the dataset includes multiple MAGs with a mean completeness score of 95%, it appears that almost all Gallionellaceae FeOB contain one of these two known mechanisms for iron oxidation. Overall, *cyc2* homologs are more common than *mtoA* (Fig. 2, Fig. 3) and some genomes encode multiple copies of *cyc2* (Table S3). All of the FeOB Gallionellaceae with *cyc2* encode at least one copy that is closely related to Cluster 1 Cyc2 (classified as in McAllister, et al. (33)), which has been functionally verified as an iron oxidase (32).

The *Ca.* Nitrotoga SL_21 MAG contains only a predicted Cluster 2 Cyc2 homolog. Confidently assigning iron oxidation function to Cluster 2 Cyc2s depends on supporting context, which is lacking in this case. *Ca.* Nitrotoga SL_21 is not from a typical iron-oxidizing environment (permanently stratified, non-marine, saline lake) and it is not closely related to the functionally verified Cluster 2 Cyc2 representative, *Acidithiobacillus*. Currently, there is no evidence that this sole *Ca.* Nitrotoga Cyc2 is an iron oxidase.

In contrast, *nxrAB* genes are exclusive to the *Ca.* Nitrotoga and copies are present in 85% of the genomes (Fig. 2, Fig. 3). Given that many of the genomes are MAGs with a mean completeness of 94%, distribution of *nxrAB* appears to indicate nitrate oxidation is the main metabolism of *Ca.* Nitrotoga. Thus, our pangenome analysis confirms Gallionellaceae can be divided into two main groups based on primary energy metabolism – FeOB and NOB.

### *c*-type cytochromes

Both known iron oxidases (Cyc2 and MtoA) in Gallionellaceae are *c*-type cytochromes that transport electrons across the outer membrane. FeOB use additional *c*-type cytochromes to transport electrons through the periplasm to the rest of the electron transport chain. We reasoned that novel iron oxidation mechanisms may also utilize *c*-type cytochromes, so we searched the Gallionellaceae genomes for proteins containing the CXXCH, CX_3_CH, and CX_4_CH heme-binding motifs (abbreviated hereafter as CXXCH). There is a stark difference between FeOB and NOB in the distribution of predicted *c*-type cytochromes. FeOB genomes have an average of 1.5x more CXXCH-containing proteins than NOB, and only non-NOB genomes encoded proteins with ten or more CXXCH motifs (Fig. 2). The abundance of genes for potential *c*-type cytochromes, in particular multiheme cytochromes (MHCs), suggest the presence of additional iron oxidation mechanisms in the Gallionellaceae FeOB.

To find *c*-type cytochromes of interest, all CXXCH-containing proteins were clustered using MMSeqs2 with bidirectional coverage and an 80% alignment cutoff. Clusters of sequences were then classified with representative sequences from isolates using BLAST to query the Uniprot database. If the cluster did not contain a sequence from an isolate, a consensus classification was used. A cluster of monoheme proteins (Cluster 313) was classified as Cyc2 and three clusters of decaheme proteins were classified as MtoA (Clusters 335 and 451) and MtrC (Cluster 50) (Fig. 2, Fig. 4, Table 1). These Cyc2, MtoA, and MtrC clusters largely agree with FeGenie’s HMM-based predicted distributions. Since MMSeqs2 generated two clusters of MtoA sequences, we sought to further verify the classifications. We constructed a tree of all Gallionellaceae MtoA sequences along with reference sequences of MtrA from iron-reducing bacteria (Fig. 5) (57). Although there is some separation of Cluster 335 and Cluster 451 MtoA sequences, many clades are not well defined or supported. In fact, backbone support throughout the tree is poor and the tree does not indicate a clear separation of the MtoA and MtrA sequences (Fig. 5).There is some evidence that the direction of electron flow through Mto/Mtr can be reversible (31, 58). So, it may be that the functions of MtoA and MtrA are interchangeable, and in fact they may be indistinguishable proteins that can conduct electrons across the outer membrane in either direction.

**FIGURE 4.**
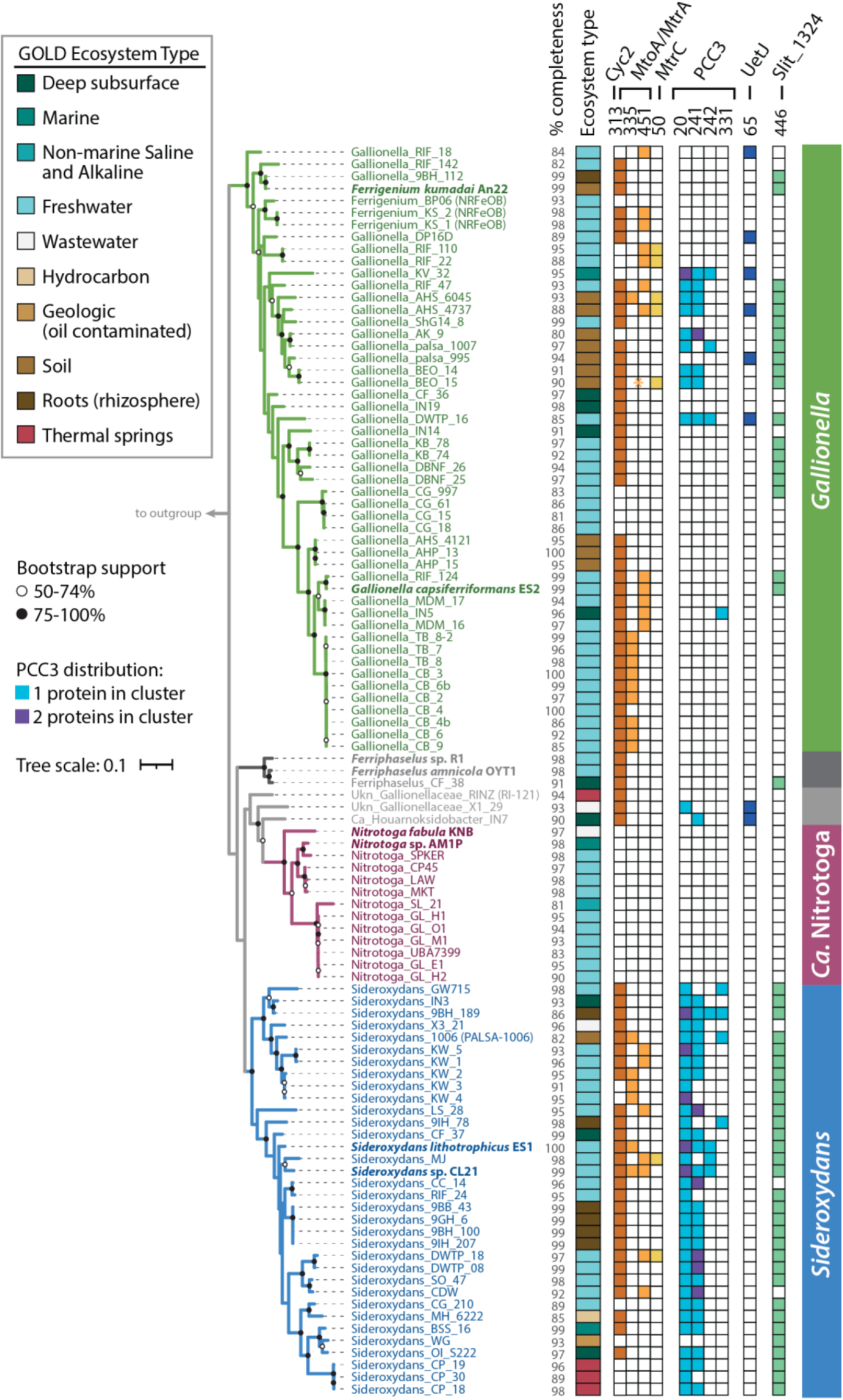
Maximum likelihood tree of concatenated ribosomal proteins from the Gallionellaceae that shows the distribution of MMSeqs2 Clusters that represent predicted cytochromes Cyc2, MtoA, MtoC, PCC3, Uet, and Slit_1324. Asterisk (*) for Gallionella_BEO_15 indicates a partial MtoA sequence was detected using HMMs and verified with BLAST, but was too short to bin into the MMseqs2 MtoA clusters. Isolates are shown in bold. % completeness = genome completeness calculated with CheckM. Outgroup omitted for space.

**TABLE 1.**
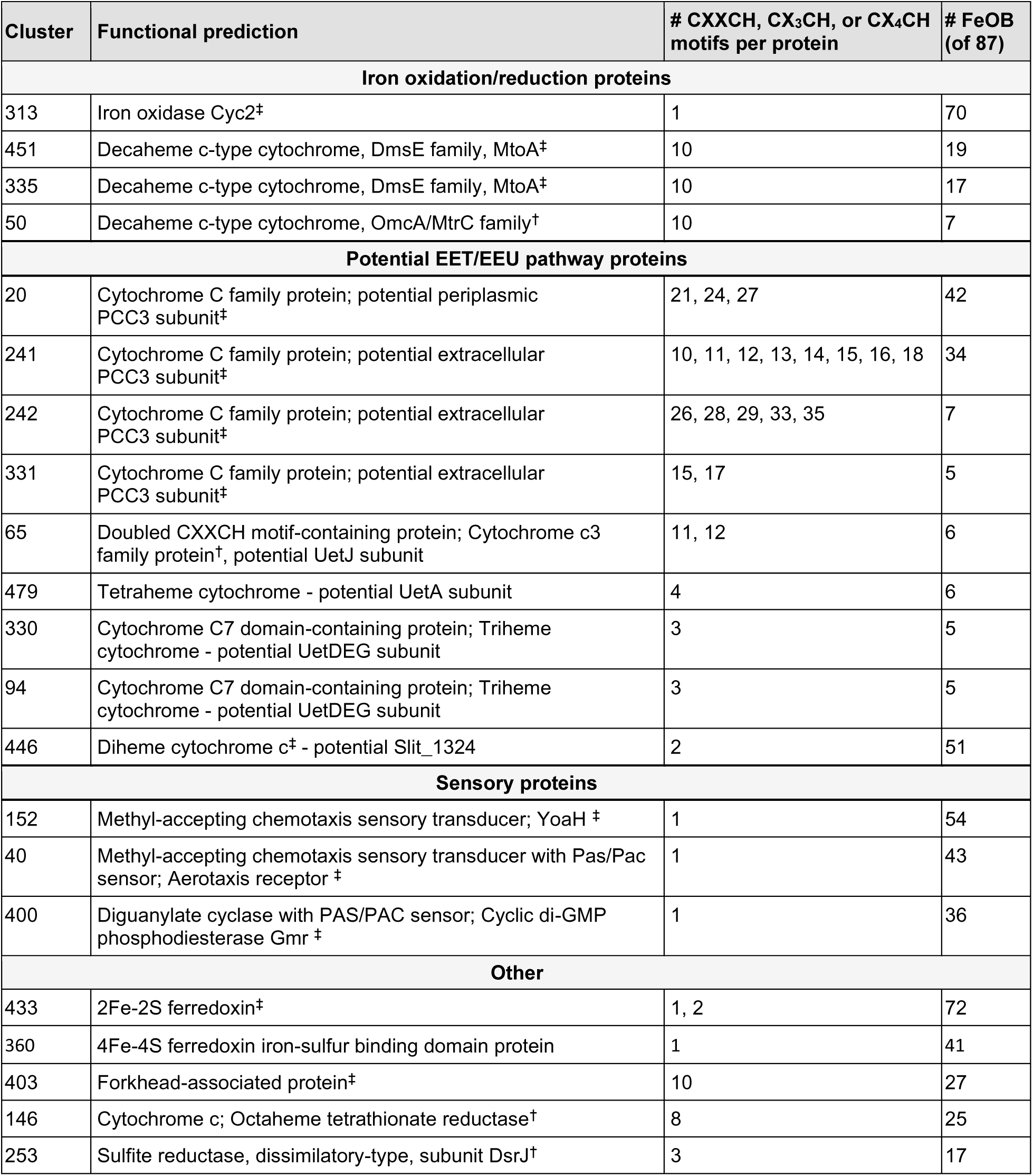
Clusters of predicted *c*-type cytochromes and other heme-containing proteins of interest from MMSeqs2. Functional predictions are based on ‡ isolate annotations and NCBI BLAST or † BLAST of sequences from metagenomes in Uniprot.

**FIGURE 5.**
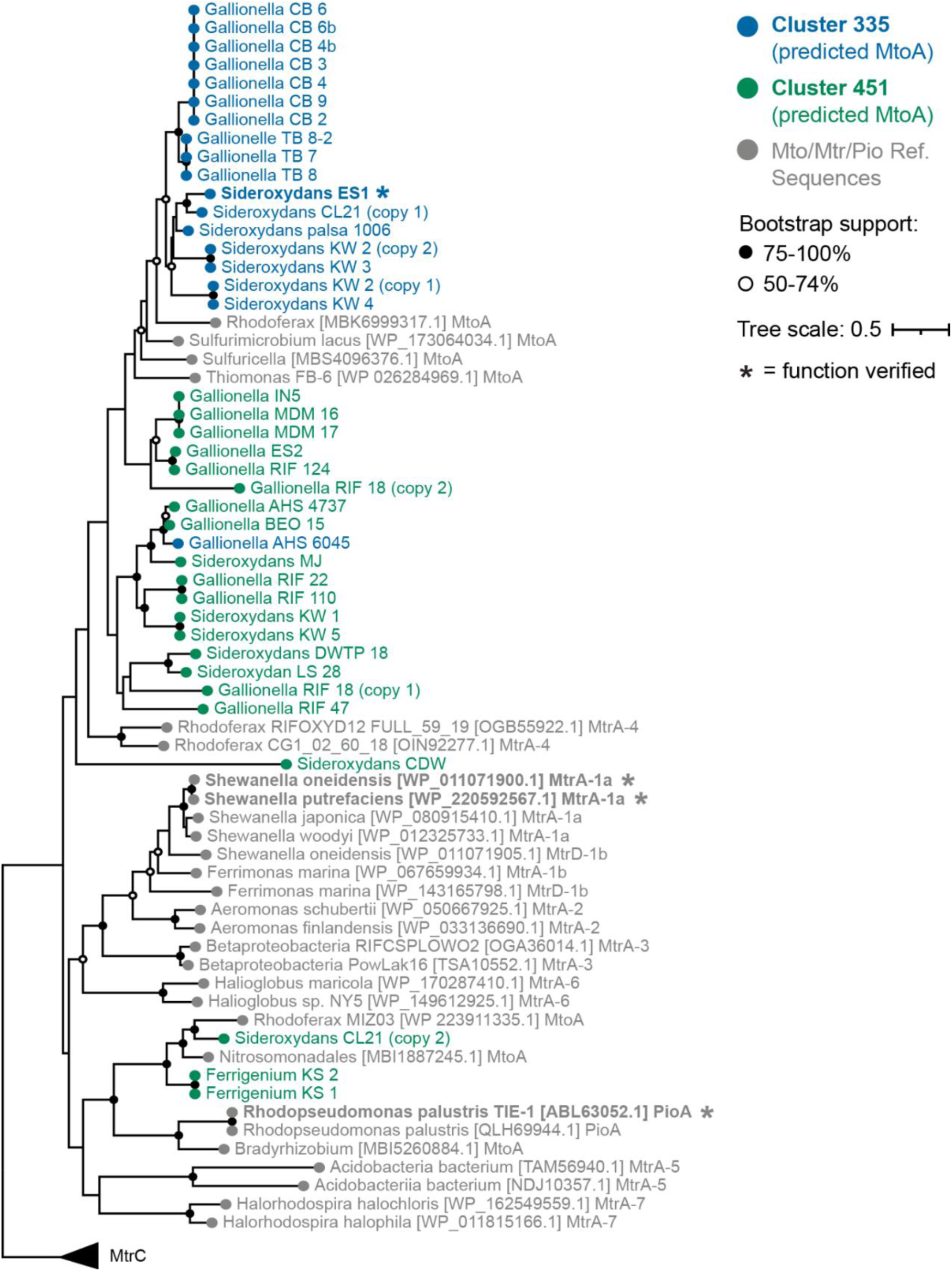
Maximum likelihood tree of the predicted MtoA sequences identified in MMSeqs2 Cluster 335 and Cluster 451 along with MtoA reference sequences from NCBI and MtrA reference sequences from Baker, et al. 2022. Numbers (1a, 1b, 2, 3, 4, 5, 6, and 7) appended after Mtr denote reference sequences from the seven MtrA groups defined by Baker, et al., 2022. MtrA-4 indicates the Group 4 Betaproteobacteria. Tree is rooted using MtrC. Support is the result of 500 bootstrap replicates.

The decaheme cytochrome MtrC is the extracellular partner of the iron-reducing MtrAB complex of *Shewanella*. The MtrAB complex is a homolog of the MtoAB complex of FeOB, but MtrC was thought to be exclusive to iron-reducers because there is no MtrC homolog in *S. lithotrophicus* ES-1. However, we found seven MAGs within both *Gallionella* and *Sideroxydans* that encode MtrC (Table S3), suggesting it may also have a role in the energy metabolism of iron oxidizers.

Multiheme porin-cytochrome *c* complexes have been proposed to play roles in extracellular electron transport and/or metal oxidation because they provide a conduit for electrons to cross the outer membrane and participate in cellular metabolism (59, 60). One example is the PCC3 complex, identified through bioinformatic analyses of genomes of several FeOB including *S. lithotrophicus* ES-1, which contains a periplasmic MHC, an extracellular MHC, an outer membrane porin, and a conserved inner membrane protein (38). We identified 26 Gallionellaceae FeOB genomes with a complete predicted PCC3 complex, an additional 11 genomes with a partial complex, and four instances where a genome’s PCC3 gene cluster encodes two predicted periplasmic cytochromes instead of one (Fig. 4, Table S3). The predicted periplasmic MHCs grouped in MMSeqs2 Cluster 20, while predicted extracellular MHCs grouped in Clusters 241, 242, and 331. The extracellular MHCs exhibited variability in the number of CXXCH heme motifs (10-35; Table 1), which suggests a range of functions in the extracellular PCC3 MHCs. Based on *in silico* protein structure models, PCC3 MHCs appear long and mostly linear (Fig. 6, Fig. S2), suggesting an extended conduction range both intra- and extracellularly.

**FIGURE 6.**
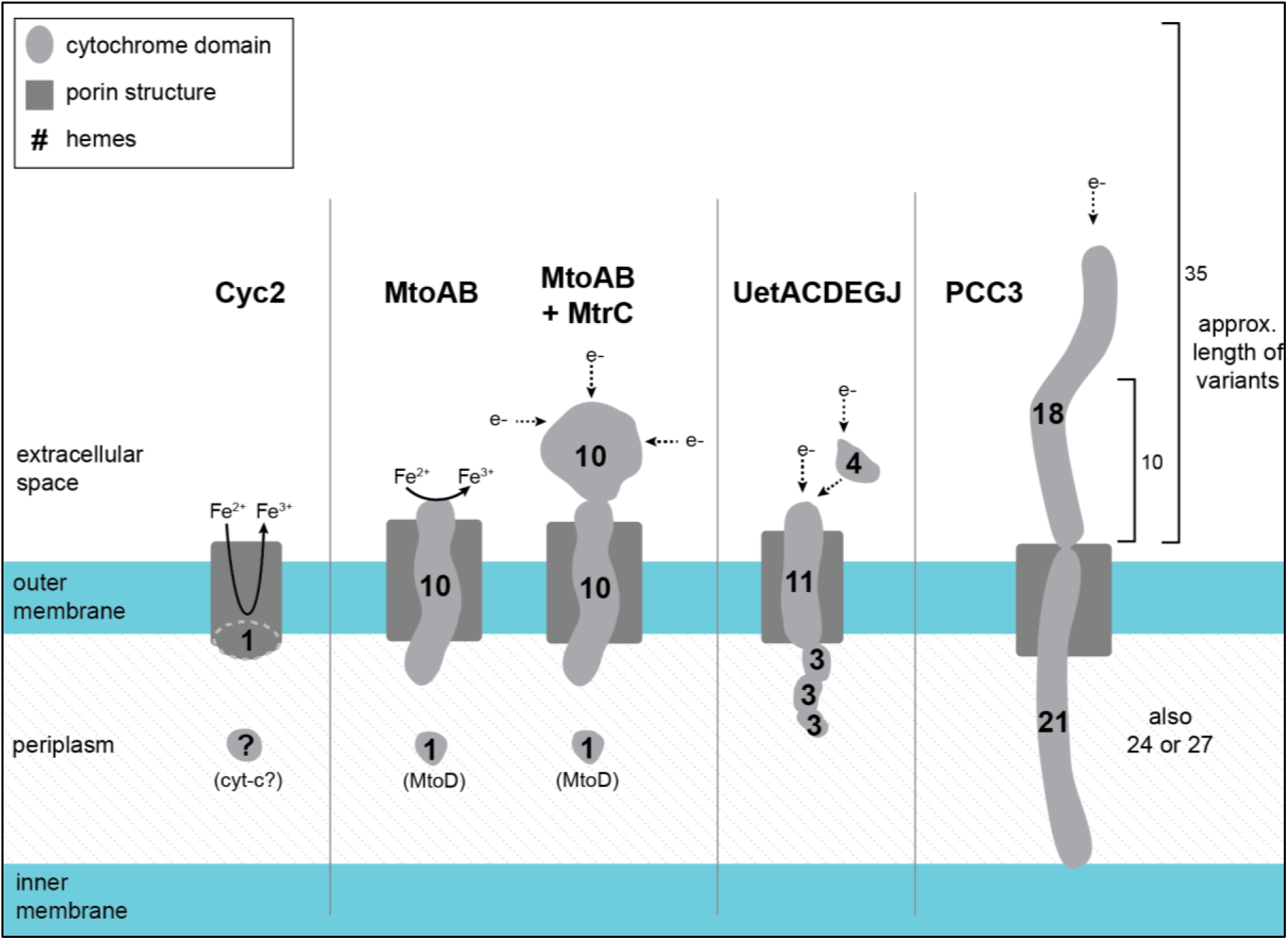
Models of potential Gallionellaceae extracellular electron transfer mechanisms. All sizes are approximated. Dimensions of Cyc2 with its fused cytochrome-porin and the porin-cytochrome complexes MtoAB, MtoAB+MtrC, MtoD and Uet drawn from models and measurements in previous literature (32, 38, 61–63). Illustration of PCC3 is based on AlphaFold2 predictions (Fig. S2). The number of hemes and size of PCC3 can vary. The 21/18 heme complex of *S. lithotrophicus* ES-1 is depicted along with the estimated length of the 10 and 35 heme variants of the extracellular cytochrome.

Another recently described multiheme porin-cytochrome *c* complex is the <u>u</u>ndecaheme <u>e</u>lectron transfer (Uet) complex, found in the cathode-oxidizing Tenderiales (61) (Fig. 6). We used a combination of MMSeqs2 and BLAST to identify Uet genes in the Gallionellaceae. While PCC3 is more common to *Sideroxydans* (59%) than *Gallionella* (12%), the Uet pathway appears exclusive to *Gallionella* and two unclassified outliers (Fig. 4). Six *Gallionella* have predicted undecaheme cytochrome (UetJ), extracellular tetraheme cytochrome (UetA), three predicted periplasmic triheme cytochromes (UetDEG), peptidylprolyl isomerase (UetB), and NHL repeat units (UetHI) (Fig. 4, Table S3). We checked for genes encoding the β-barrel porin UetC and found BLAST hits in four of the six genomes (Table S3).

*S. lithotrophicus* ES-1 has a set of periplasmic cytochrome genes without a predicted porin that were highly upregulated during growth on iron, and therefore thought to be involved in iron oxidation (27). The genes encode a cytochrome b (Slit_1321), a hypothetical extracellular protein (Slit_1322), a monoheme cytochrome class I (Slit_1323), a periplasmic diheme cytochrome (Slit_1324; Cluster 446 in Table 1), and a molecular chaperone Hsp33 (Slit_1325). We found homologs of the Slit_1321-1324 gene cluster are common and well-conserved among Gallionellaceae FeOB, present in 50 genomes (Fig. 4, Table S3). These genes may represent a mechanism of periplasmic electron transport, perhaps as part of an iron oxidation/extracellular electron uptake pathway.

### Electron transport chains

We compared electron transport chain component genes of the iron and nitrite oxidizer groups and found them to be largely similar (Fig. 7). High-affinity *cbb*_3_-type oxidases are common (Fig. 3), with most genomes containing either the proximal or distal form of *ccoN* (Fig. 2) (64). Even the four NRFeOB genomes contain *ccoNO* genes, indicating a potential for both oxygen and nitrate respiration. In contrast, few Gallionellaceae genomes contain *narGH* or *napAB* (6 and 10 genomes, respectively, with no overlap), indicating nitrate respiration is relatively rare overall (Fig. 3, Table S3).

**FIGURE 7.**
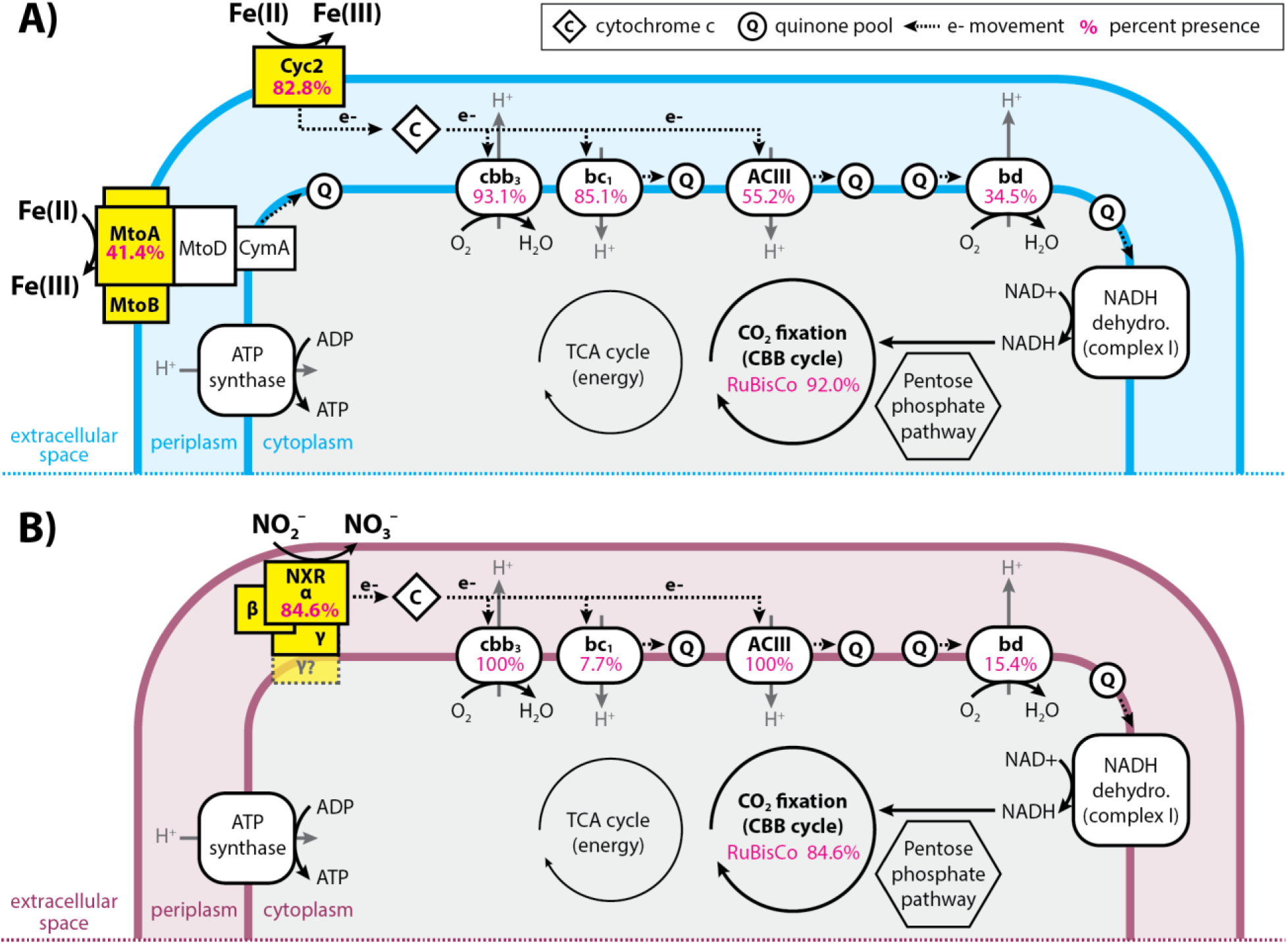
Diagram showing the similarities and differences between the electron transport chains of (A) iron-versus (B) nitrite-oxidizing Gallionellaceae. Pink numbers indicate the percent of FeOB (A) or NOB (B) genomes that encoded each part of the electron transport chain or RuBisCo.

In addition to the *cbb*_3_-type oxidase genes, 34.5% of iron oxidizers and 15.4% of nitrite oxidizers possess genes for cytochrome *bd*-type oxidases (*cydAB*) (Fig. 7). The presence of *bd*-type oxidase genes often overlaps with *cbb*_3_-type oxidase genes (Table S3). Like *cbb*_3_-type oxidases, cytochrome *bd*-type oxidases have a high affinity for oxygen and recent studies show they can be more highly expressed than *cbb*_3_-type oxidases under low-oxygen, organic-rich conditions (65). Both FeOB and NOB have genes for cytochrome *bc*_1_ and Alternative complex III (ACIII) quinol oxidase complexes. However, *bc*_1_ is more common in FeOB (85.1%) compared to NOB (7.7%), while ACIII is more common in NOB (100%) than FeOB (55.2%) (Fig. 7, Table S3). Like the *bd*- and *cbb*_3_-type oxidases, presence of *bc*_1_ and ACIII often overlaps in a single organism, especially in FeOB (Table S3). Possessing both *bd*- and *cbb*_3_-type oxidases, and/or having both *bc*_1_ and ACIII contributes to flexibility within the electron transport chains of Gallionellaceae. The presence of various terminal oxidases implies adaptation to niches where oxygen and organic carbon availability differ or fluctuate.

### Carbon fixation

Gallionellaceae isolates grow autotrophically. To determine if the capacity for autotrophic growth is widespread, we analyzed the pangenome for RuBisCo genes (*cbbLS*, *cbbMQ*). Most genomes in the dataset (>91%; 94 of 103 genomes) contain genes for either Form I or Form II RuBisCo (Fig. 3, Table S3). FeOB more commonly have Form II, while NOB only have Form I. Form II enzymes are adapted for medium to high CO_2_ and low O_2_ concentrations (66) and their predominance in FeOB may correspond to different oxygen niches of FeOB and NOB. The prevalence of RuBisCo genes indicates both iron- and nitrite-oxidizing Gallionellaceae have the capacity to grow autotrophically.

### Auxiliary energy metabolisms

Previous studies showed some Gallionellaceae FeOB possess alternate energy metabolisms such as thiosulfate and lactate oxidation (27, 28). We searched the pangenome for key genes of sulfur, manganese, and organic substrate oxidation pathways to determine how common alternate metabolisms are among Gallionellaceae FeOB. Sulfide:quinone reductase (*sqr*) is common to both FeOB and NOB (Fig. 3, Table S3). Sqr can oxidize sulfide, transporting electrons to the quinone pool, although it may be a means of detoxification rather than energy conservation (67, 68). In contrast, both *soxABXYZ* and *dsrAB* are detected exclusively in the iron-oxidizing Gallionellaceae genomes (Fig. 3, Table S3). To predict the oxidative vs. reductive function of *dsrAB,* we constructed a tree using reference sequences from Loy, et al. (69, 70). Gallionellaceae sequences form a discrete clade within the sulfur-oxidizing group (Fig. S3), indicating the DsrAB of Gallionellaceae is likely a reverse dissimilatory sulfite reductase (rDSR). In contrast, the *Ca.* Nitrotoga genomes do not contain *dsr* or *sox* genes. Instead, *Ca.* Nitrotoga have *sorAB*, which may enable oxidation of sulfite to sulfate (Fig. 3). Together these results indicate that although sulfur oxidation is an accessory trait of both iron- and nitrite-oxidizing Gallionellaceae, only certain FeOB appear capable of oxidizing S(0) or thiosulfate.

We analyzed the pangenome for signs of organic utilization. Although not widely distributed, the most common genes were for lactate utilization (*lutABCP*) and sugar transport (*msmX, gtsABC*). Only eight *Gallionella* and five *Sideroxydans* genomes, including *Sideroxydans* sp. CL21, have *lutABC* along with the *lutP* lactate permease gene (Fig. 2, Fig. 3, Table S3). Likewise, only six genomes contain *gtsABC* genes for glucose/mannose uptake. None of the NOB contain the *lut* or *gts* genes for organic utilization.

We used BLAST to evaluate the Gallionellaceae genomes for manganese oxidase genes *mcoA*, *moxA*, *mofA*, and *mnxG*. There are a few hits for *mcoA*, *moxA*, and *mofA* genes, but none for *mnxG* (Table S3). Since manganese oxidation activity has not been shown in any of the Gallionellaceae isolates, additional verification is needed to determine whether the genes identified by BLAST are truly Mn oxidases.

### Other genes distinct to FeOB, potentially related to iron oxidation

We searched the pangenome for the candidate genes for stalk formation (*sfz/sfb*) identified in the stalk-forming *Ferriphaselus* and Zetaproteobacteria isolates (71, 72). Twelve genomes, restricted to *Gallionella* (9) and *Ferriphaselus* (3) contain the four *sfz/sfb* genes (Fig. 2, Table S3). Thus far, all cultured Gallionellaceae stalk formers belong to these two genera, suggesting stalk formation may be limited and not a trait of *Sideroxydans*.

Using the Anvi’o subset of only genomes >97% complete, we identified several gene clusters that were present and abundant only in *Gallionella* and *Sideroxydans*, but lacked prior connection to an iron-oxidizing lifestyle. These included distinct gene clusters with COG functional annotations for: Cell Wall/Membrane/Envelope Biogenesis, Cytoskeleton formation, Signal Transduction Mechanisms, and Energy Production and Conversion (Table 2, Table S4). Clusters for Cell Wall/Membrane/Envelope Biogenesis may indicate FeOB have specific adaptations for housing *c*-type cytochromes and EET mechanisms in the outer membrane, or to avoid encrustation by iron oxides. Clusters for Energy Production and Conversion included ferredoxin (Fdx) and subunits of the RnfABCDEG complex. The Rnf complex was originally discovered for its role in N fixation, in which it oxidizes NADH and generates reduced ferredoxin that donates electrons to nitrogenase (73). More recent studies have shown Rnf complexes can conserve energy under anaerobic conditions (74–76) and, as a low potential electron donor, ferredoxin can transfer electrons to many metabolic pathways including some that produce secondary metabolites (77). Not all Gallionellaceae with Rnf complex genes have *nifDHK* nitrogenase genes, implying Gallionellaceae Rnf and ferredoxin have functions beyond N fixation. Although their specific function in Gallionellaceae FeOB are unknown, their ubiquity implies utility for FeOB and an area for additional research.

**TABLE 2.**
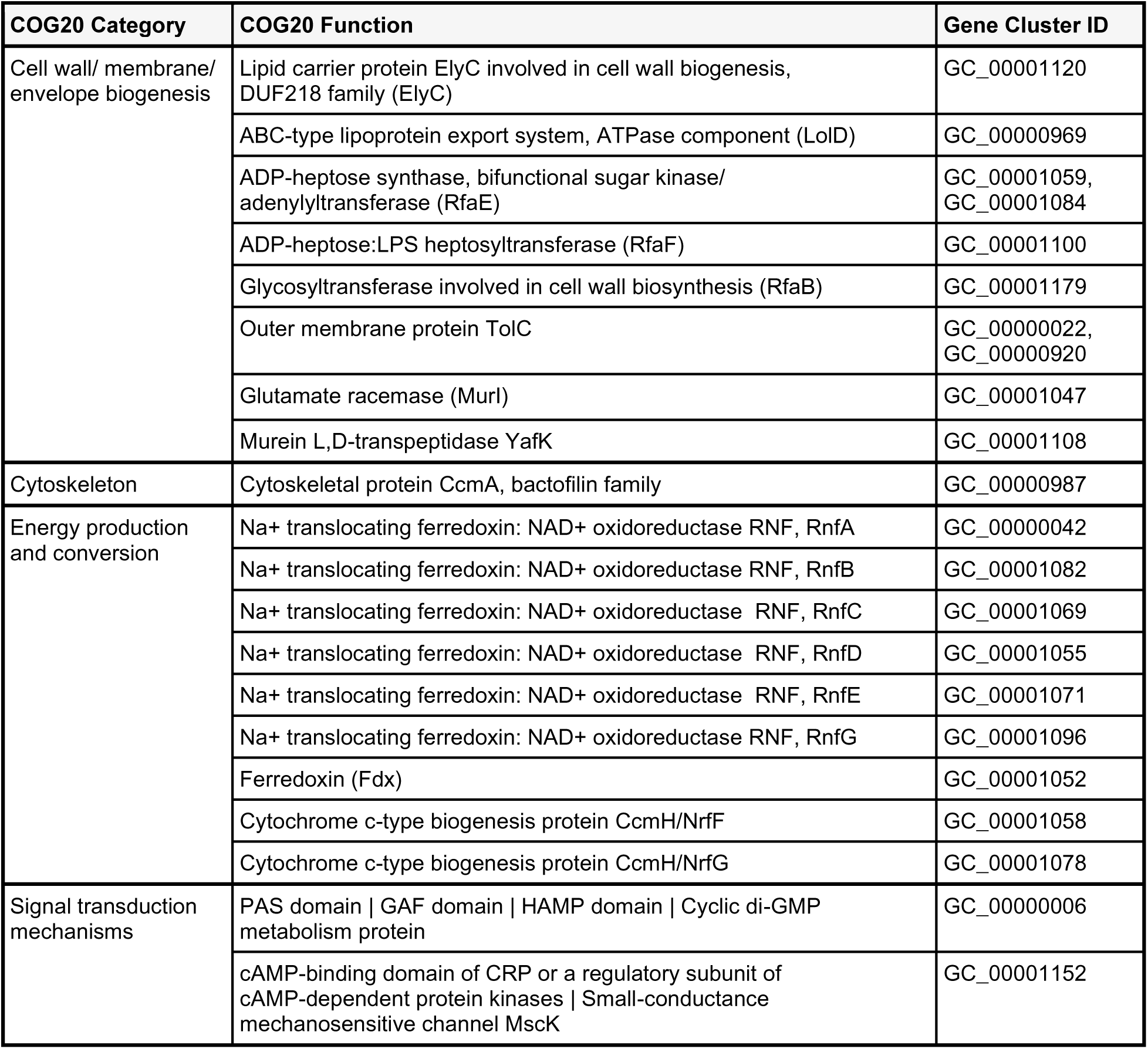
Gene clusters of interest from the Anvi’o pangenome subset that were present in iron-oxidizing *Gallionella* and *Sideroxydans*, but absent in nitrite-oxidizing *Ca.* Nitrotoga.

## Discussion

The Gallionellaceae family is historically known for its iron-oxidizing members, but recently a new candidate genus of nitrite oxidizers, *Ca.* Nitrotoga, was identified (39).

Comparing their genomes to those of FeOB genera has helped identify genes and pathways related to iron oxidation since *Ca.* Nitrotoga isolates have no documented capacity for that metabolism (39, 40, 42, 43, 45). We resolved the phylogeny of the Gallionellaceae and verified *Ca.* Nitrotoga lacked iron oxidation marker genes. Given separate groups of FeOB and NOB, we used a pangenomic approach to identify shared features of the Gallionellaceae, as well as FeOB-specific genes that may represent novel iron oxidation pathways.

### Phylogeny

The Gallionellaceae is composed of four genera: *Gallionella, Sideroxydans*, *Ferriphaselus*, and *Ca.* Nitrotoga, based on a concatenated ribosomal protein tree. In comparison, 16S rRNA phylogeny did a poorer job of resolving these genera, so 16S-based identification should be considered tentative, pending availability of genomes. To facilitate consistent classification, the protein sequences and alignments used here (Fig. 1) are available at (https://figshare.com/projects/Gallionellaceae_Ribosomal_Proteins_for_Concatenated_Tree/157 347).

The new phylogeny provides a framework for understanding the diversity and major metabolisms of the Gallionellaceae. They are members of Nitrosomonadales, which contain many chemolithotrophic S and N oxidizers. Like their closest relatives, the Sulfuricellaceae (78), many Gallionellaceae retain the ability to oxidize sulfur (Fig. 3, Fig. S3). The Gallionellaceae tree (Fig. 1) shows a deeply branching split between genera, with each of the two major genera, *Gallionella* and *Sideroxydans*, containing a continuum of diversity. Within the *Gallionella*, the isolates *G. capsiferriformans* ES-2 and *Ferrigenium kumadai* An22 bracket the *Gallionella*, with An22 deeply branching and ES-2 at the crown. *F. kumadai* An22 was originally classified as *Ferrigenium* based on 16S rRNA distance (25). However, our analyses do not show any clear phylogenetic clustering or functional distinction, with which we could draw a line between *Gallionella* and *Ferrigenium*. Moreover, the tree topology suggests continued diversification within both *Gallionella* and *Sideroxydans* largely without the formation of subclades that represent distinct niches. There is one subclade of *Sideroxydans* that corresponds to the GTDB genus level designation PALSA-1006 (Fig. 1). However, ANI/AAI results (Table S5) indicate there is not enough diversity within the Gallionellaceae to justify further splitting the four major genera any further. Additionally, we did not detect any obvious functional difference in PALSA-1006. Given our phylogenetic analysis, ANI/AAI, and similar functional profiles, we recommend keeping them within *Sideroxydans*. Based on the above classification scheme, most of the genomes (84 of 103) fall into either *Gallionella* or *Sideroxydans*.

Phylogenetic diversity corresponds to functional diversity that can drive Gallionellaceae success in a variety of environments. Many *Gallionella* and *Sideroxydans* do not appear to be obligate iron oxidizers, and some may not be obligate aerobes. Auxiliary metabolisms for S, N, and C are present to varying degrees throughout the iron-oxidizing genera and are not associated with specific sub-groups. Some FeOB from organic-rich environments, such as *Sideroxydans* sp. CL21, have genes for organoheterotrophy. Other FeOB show metabolic flexibility in additional lithotrophic metabolisms, such as oxidation of S or potentially Mn, elements that often co-occur with Fe in the environment. Some Gallionellaceae may also thrive in oxygen-poor environments by reducing nitrate, although this capability appears rare. Such traits contribute to diversity in the Gallionellaceae FeOB genera, which appear to acquire and/or retain additional energy and nutrient metabolisms to adapt to a range of environments.

*Ca.* Nitrotoga stands out as an exception within the Gallionellaceae. The pangenome analysis shows that *Ca.* Nitrotoga have distinctive genomic content (Fig. S4). They do not appear to have the capacity for iron oxidation based on available physiological evidence and the genomic analyses presented here. The similarities in Gallionellaceae FeOB and *Ca*. Nitrotoga electron transport chains enable them to meet the shared challenge of conserving energy from high-potential electron donors. However, *Ca*. Nitrotoga are a distinct clade that appear to have evolved from the FeOB to occupy a nitrite oxidation niche.

### Iron oxidation and extracellular electron uptake mechanisms

The Gallionellaceae FeOB genomes encode a wide variety of predicted *c*-type cytochromes. Of these cytochromes, many appear to be associated with the outer membrane, implying a role in extracellular electron transport. Cyc2 is present in the majority of Gallionellaceae FeOB genomes, while multiheme cytochromes (MHC) Mto/Mtr, Uet, and PCC3 are less common, each with different distribution patterns (Fig. 4), suggesting the different cytochromes play distinct roles.

Cyc2 has been shown to oxidize dissolved Fe(II) (27, 32, 37). The monoheme Cyc2 is a small fused cytochrome-porin and since aqueous Fe^2+^ is common to many redox transition zones, it makes sense that most FeOB would retain and use the simplest tool. But in Earth’s various environments, iron is largely available as minerals (clays, oxides, sulfides) and also bound to organics (e.g. humic substances). The decaheme MtoA has been shown to play roles in the oxidation of mineral-bound Fe(II), specifically Fe(II) smectite clay (37). As a MHC, MtoA may have multiple benefits that help in oxidizing minerals. MtoA has a large redox potential window (−350 to +30 mV; (31, 60)), which could help with oxidation of solids, like smectite, that also have a range of redox potentials (e.g., −600 to +0 mV for SWa-1 vs. −400 to +400 mV for SWy-2; (79)), which change as mineral-bound iron is oxidized or reduced. Assuming the MtoA structure is similar to MtrA, the ten hemes span the membrane, making a wire that conducts from extracellular substrates to periplasmic proteins (62, 80). The multiple hemes allow for transfer of multiple electrons at a time (59). Some MAGs with *mtoAB* also encode the extracellular decaheme cytochrome MtrC. In *Shewanella,* the MtrCAB complex requires MtrC to reduce solid minerals (ferrihydrite), while MtrAB alone can only reduce dissolved Fe(III) and electrodes (81–83). Likewise, Gallionellaceae MtrC may help increase interactions with different minerals. Some Gallionellaceae FeOB may retain genes for both Cyc2 and MtoAB (with or without MtrC) to oxidize different Fe(II) substrates in their environments.

Like MtrCAB, the predicted PCC3 complex includes both periplasmic and extracellular MHCs and a porin. A key difference is that the PCC3 cytochromes often have more hemes than MtoA/MtrA and MtrC. The greater number of hemes may serve to store electrons, as in a capacitor. They may also conduct across a greater distance; the PCC3 periplasmic MHC, with 21-27 hemes, is potentially long enough to span the entire periplasm (as noted by Edwards et al., (84)). Intraprotein electron transfer between hemes is rapid (85–87); therefore the periplasm-spanning MHC of PCC3 may allow for faster electron transfer compared to complexes containing smaller periplasmic cytochromes like the monoheme MtoD. The extracellular PCC3 MHC contains between 10 and 35 hemes, which could extend further from the outer membrane compared to MtrC. Not only would this extend the range of electron transfer, but may also be faster than a “wire” of smaller cytochromes (e.g. *Geobacter* hexaheme OmcS (88)). Increasing oxidation rates via larger MHCs would allow FeOB to oxidize substrates faster. Given that Fe(II) is subject to abiotic oxidation under certain conditions and other organisms may compete for EEU, such kinetic advantages would give FeOB a competitive edge.

## Conclusions

Gallionellaceae, specifically *Gallionella*, is best known for lithoautotrophically oxidizing iron to make mineral stalks that come together to form microbial mats at groundwater seeps (18, 89, 90). Although this may contribute to an impression that the niche was relatively restricted, 16S rRNA sequencing of cultures and environmental samples has revealed both the diversity of Gallionellaceae as well as its prevalence across practically any freshwater and some brackish environments where Fe(II) and O_2_ meet. The pangenome shows that Gallionellaceae possess metabolic flexibility to use non-iron substrates, notably sulfur, and the MHCs likely also confer further metabolic capabilities that may help them occupy a range of different iron- and mineral-rich niches. Gallionellaceae thrive in aquifers, soil, and wetlands, all of which have substantial mineral content. Thus, the widespread ecological success of Gallionellaceae may well correspond to genomes that encode a range of iron oxidation mechanisms as well as adaptations for varied environments.

It is becoming clear that there are multiple ways to oxidize iron, though we have varying levels of evidence for gene/protein function (60, 91, 92). Validating iron oxidation genes/proteins is painstaking work due to challenging cultures, low yield, few genetic systems, and the fact that iron interferes with many molecular extractions and assays. And yet, there are likely even more iron oxidation mechanisms, so we need to be strategic about choosing genes/proteins for deeper characterization. Our pangenome analysis gives a wider view of the distribution and frequency of potentially novel iron oxidation genes, which will help us to prioritize investigations. Furthermore, the varied outer membrane-associated cytochromes inspire us to investigate relationships between structure and function. Why are there so many different multiheme cytochromes? Is there substrate specificity, kinetic advantages, battery-like functions, or some utility we have yet to consider? Addressing these questions will help us understand how these proteins and pathways shape microbial transformations of varied Earth materials.

## Methods

### Data collection and curation

Gallionellaceae genomes were collected from the National Center for Biotechnology Information (NCBI) Entrez database (93), the Joint Genome Institute Integrated Microbial Genomes (JGI IMG) database (94), and the European Nucleotide Archive (ENA) at EMBL-EBI database (*Sideroxydans* sp. CL21, *Ca.* Nitrotoga fabula KNB, and the “IN” MAGs (17, 43, 95)) (Table S6). We also received non-public genomes from the Ménez Lab at the Université de Paris (3 genomes reconstructed by Aurélien Lecoeuvre from the Carbfix study in Hengill, Iceland (2); metagenomes available at Sequence Read Archive SRR3731039, SRR3731040, SRR4188484, and SRR4188643), and the Banfield Lab at the University of California, Berkeley (3 genomes reconstructed by Alex Probst from Crystal Geyser in Utah, USA (96)) (Table S6). This initial 230-genome dataset included isolate genomes, metagenome assembled genomes (MAGs), and single-cell amplified genomes (SAGs) that were taxonomically classified as members of the Gallionellales order; Gallionellaceae family; or the *Gallionella, Sideroxydans*, *Ferriphaselus*, *Ferrigenium*, or *Ca.* Nitrotoga genera in their respective databases. Duplicate genomes were identified and removed if they had identical accession numbers or their average nucleotide identities (ANI) were 100%. CheckM v1.1.2 (97) was used to assess genome quality. Genomes with lower than 80% completeness and greater than 7% contamination were removed from the dataset. The final filtered dataset, referred to as “the Gallionellaceae” or “the dataset” contained 103 genomes, including six of the Gallionellaceae FeOB isolates (Table S1). The seventh isolate, *Sideroxyarcus emersonii* (26), was not published at the time of our main analysis, but a supplemental of its key metabolic genes and MHCs (Table S7) shows it has similar patterns to *Sideroxydans*.

#### Naming conventions

To assign simple, unique names to the metagenomes, codes were appended to genus-level names based on sample location and bin IDs (Table S1, Table S6, Table S8). Isolates retained their own unique names. Organisms that were taxonomically classified in their original databases at the family Gallionellaceae or order Gallionellales were, if possible, classified at lower taxonomic levels using a combination of AAI, 16S rRNA (if available), classification through the Genome Taxonomy Database Toolkit (GTDB-Tk) (98), and placement in the concatenated ribosomal protein tree (Fig. 1 and Fig. 2).

#### Ecosystem classifications

To assess whether metabolic diversity correlated to ecosystem type, each genome was assigned to an ecosystem based on the Genomes OnLine Database (GOLD) (99) schema which leverages Environmental Ontology (EnvO) classifications (100). A genome’s pre-existing classification from IMG was used if available. Genomes without prior classification were categorized based on published descriptions of their sample sites and “habitat” information listed in their database of origin. Based on the GOLD classifications (Table S2), genomes were examined for patterns of correspondence between ecosystems and phylogenetic and/or metabolic diversity.

### Calculation of average amino acid and nucleotide identities

Average amino acid identity (AAI) and average nucleotide identities (ANI) were computed to assess the similarity of genomes in the curated data set (Table S5). AAI was calculated using CompareM (101). ANI was calculated using FastANI in Kbase (102). Final AAI and ANI tables were formatted using Microsoft Excel.

### Tree construction

#### Concatenated ribosomal protein tree

A concatenated tree of ribosomal proteins (Fig. 1) was constructed to determine the phylogenetic relationships of genomes in the Gallionellaceae dataset. Two *Sulfuricella* genomes, *Sulfuricella* sp. T08 and *Sulfuricella* 3300027815, where included as an outgroup to root the tree. Use of a *Sulfuricella* outgroup was based on previous literature (103, 104), which identified *Sulfuricella* and other members of the Sulfuricellaceae family as near neighbors of Gallionellaceae. The concatenated sequences were composed of 13 small and large ribosomal proteins (L19, L20, L28, L17, L9_C, S16, L21p, L27, L35p, S11, S20p, S6, S9) present in 94 or more of the 105 genomes including the outgroup. Protein sequences were aligned in Geneious v.10.2.6 (105) using MUSCLE (106). Ends of the alignments were manually trimmed and regions with over 70% gaps were masked, after which sequences were concatenated. The tree was constructed using RAxML-NG v1.0.3 (107) with the maximum likelihood method, LG+G model, and 1000 bootstraps. The final tree was visualized and annotated with iTOL (108).

### 16S rRNA gene tree

We constructed a 16S rRNA gene tree (Fig. S1) composed of sequences from our dataset combined with a selection of sequences from the SILVA database to determine how well 16S rRNA resolves Gallionellaceae phylogeny compared to the concatenated ribosomal protein tree. Full length (∼1500 bp) 16S rRNA genes were retrieved from 24 of the Gallionellaceae genomes using Anvio’s ‘anvi-get-sequences-for-hmm-hits’ command for “Ribosomal_RNA_16S.” These genes were aligned in SINA (109) along with Gallionellaceae sequences from the Silva database (110) that had >1475 bp and >85-90 sequence quality score. The outgroup is composed of *Thiobacillus*, *Ferritrophicum*, *Sulfuricella*, *Sulfuriferula*, and *Nitrosomonas* sequences acquired from the Silva database. The final alignment contained 965 non-redundant sequences and alignment length was 1500 positions after trimming and masking all sequence gaps greater than 70%. A maximum likelihood tree was constructed using RAxML-NG v1.0.3 (107) with the GTR+G model and 300 bootstraps. Family and genus level classifications from the SILVA database were used to annotate the tree in Iroki (111).

#### Individual protein trees

Trees for DsrAB (Fig. S3) and Mto/Mtr (Fig. 5) were constructed from Gallionellaceae protein sequences along with reference sequences from NCBI, Loy, et al. and Baker, et al. (57, 69). Sequences were aligned with MUSCLE (106), ends were manually trimmed, and regions with over 70% sequence gaps were masked in Geneious v.10.2.6 (105). For the Dsr tree, DsrA and DsrB sequences were concatenated. Trees were constructed using RAxML-NG v1.0.3 (107) with the LG+G model. Branch support for Mto/Mtr tree is based on 500 bootstraps and support for the DsrAB tree is based on 300 bootstraps. Final trees were visualized and annotated with Iroki (111).

### Pangenome analysis

#### Metabolic gene analysis

We used the Distilled and Refined Annotation of Metabolism (DRAM) v0.0.2 (48) within KBase (102), LithoGenie within MagicLamp (50), and FeGenie (49) to identify key metabolic genes indicative of various oxidation, respiration, and carbon utilization pathways. NCBI BLAST+ (53) was used to identify additional genes for eNOR, cNOR, SorAB, Mn oxidases, LutABCP, and stalk formation. We then analyzed the presence/absence of the metabolic genes and looked for patterns across the concatenated protein tree, between genera, and between FeOB versus NOB.

#### Multiheme cytochrome analysis

To identify potential *c*-type cytochromes we used a modified heme counter script (54) to search for CXXCH, CXXXCH and CXXXXCH motifs within the protein sequences of each genome. The search identified 5,929 protein sequences with one or more CX_2-4_CH-motifs. To determine which protein sequences were shared between genomes, sequences were clustered using MMSeqs2 (112) with coverage mode 0 for bidirectional coverage of at least 80% of the query and target sequences. Several clusters of interest were identified based on either the number of CX_2-4_CH-motifs in each sequence or the relative abundance of FeOB sequences in the cluster. Querying with BLASTp (52) against the Uniprot (113) database was used to classify sequences from clusters of interest thereby identifying clusters of predicted *c*-type cytochromes. Isolate sequences were used as representative sequences for cluster classification. If a cluster did not contain an isolate sequence, a consensus classification was used. The subcellular localization of proteins was predicted using a combination of PSORTb v3.0.3 (114) and LocTree3 (115).

Some MHCs were predicted to be part of Mto, PCC3, or Uet porin-cytochrome complexes. Therefore, we wanted to determine if the genes for these MHCs were colocalized in their respective genomes with genes for β-barrel porins, periplasmic proteins, and inner membrane proteins previously identified in the literature (38, 61). We searched for the associated genes using BLASTp and amino acid reference sequences from *S. lithotrophicus* ES-1 (MtoB, MtoD, CymA), *Gallionella* AHS-4737 (MtoC), and *Ca.* Tenderia electrophaga (UetBCDEFGHI). The locus tags of BLASTp hits were then compared to locus tags of the MHCs to evaluate synteny and colocalization. The same method was used to determine if diheme *c*-type cytochromes from MMseqs2 cluster 446 which includes Slit_1324 were colocalized with a cytochrome b (Slit_1321), hypothetical extracellular protein (Slit_1322), monoheme cytochrome class I (Slit_1323), and molecular chaperone Hsp33 (Slit_1325).

#### PCC3 modeling

To model predicted PCC3 proteins, we used ColabFold: AlphaFold2 using MMseqs2 (116). Setting included using MSA mode “MMseqs2 (UniRef+environmental),” pair mode “unpaired+paired,” protein structure prediction with “AlphaFold2-ptm,” and complex prediction with “AlphaFold-multimer-v2” (117, 118). The best scoring model was rendered in PyMol v2.5.4 (119).

#### Anvi’o subset analysis

We used the Anvi’o v7 (54, 56) to build a pangenome database of all *Gallionella* (16), *Sideroxydans* (15), and *Ca.* Nitrotoga (6) genomes that were over 97% complete (Fig. S4) to analyze for additional genes important to FeOB lifestyles. Genes were clustered within the Anvi’o pangenome using a min-bit parameter of 0.5 and an mcl inflation parameter of 2. The Anvi’o pangenome was used to compare gene clusters across the dataset and to bin: 1) near-core (found in >85% of genomes), 2) accessory (found in >1 but <85% of genomes), and 3) strain specific (found in a single genome) sets of gene clusters. Gene annotations were assigned in Anvi’o using Prodigal (120) and functional annotations for Anvi’o gene clusters were assigned using the NCBI’s Database of Clusters of Orthologous Genes (COGs) (121, 122). Data tables of the binned Anvi’o gene clusters were analyzed to identify gene clusters found in the near-core genomes of *Gallionella* and *Sideroxydans* but absent in *Ca.* Nitrotoga.

## Acknowledgements

This research was funded by the National Science Foundation (EAR-1833525 to C.S.C. and S.W.P., MCB-1817651 to C.S.C.) and the Office of Naval Research (N00014-17-1-2640 C.S.C.). R. H. was also supported by fellowships from University of Delaware Graduate College and the Microbiology Program/Unidel Foundation. Support from the University of Delaware Center for Bioinformatics and Computational Biology Core Facility (RRID:SCR_017696) including use of the BIOMIX compute cluster was made possible through funding from Delaware INBRE (NIH P20GM103446), the State of Delaware, and the Delaware Biotechnology Institute.

We thank Aurélien Lecoeuvre, Emmanuelle Gérard, Bénédicte Ménez, Alex Probst, and Jill Banfield for allowing us to use unpublished genomes from their private collections; and all those who granted us permission to use their publically available, unpublished genomes from the NCBI and IMG databases (Table S1, Table S6).

We declare that we have no conflicts of interest.

