## Supplemental Figures for "Gallionellaceae pangenomic analysis reveals insight into phylogeny, metabolic flexibility, and iron oxidation mechanisms"

Number of pages: 8

Number of figures: 4

Number of tables: 7

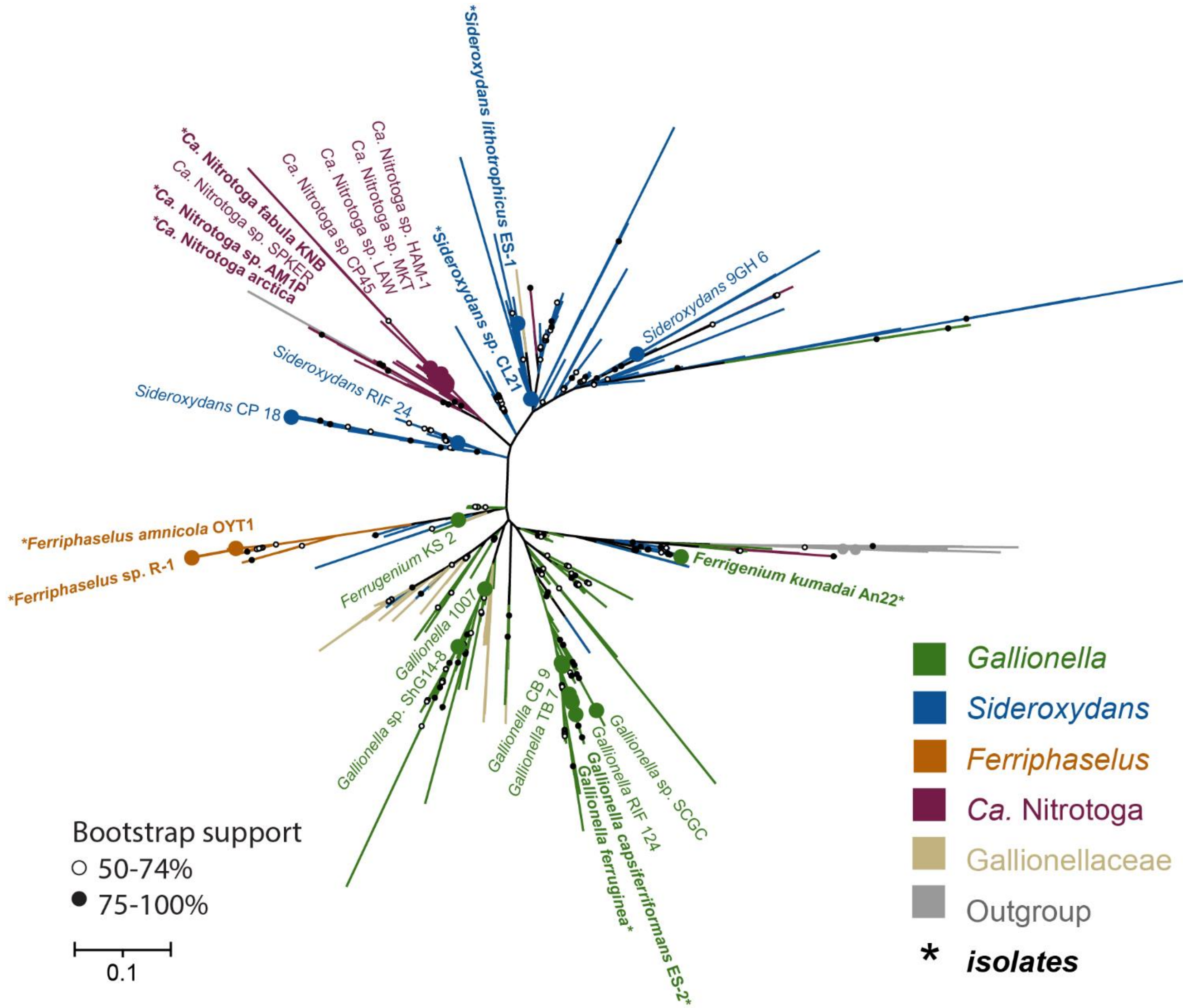


**FIGURE S1** 16S rRNA gene tree of the Gallionellaceae in this study along with 941 Gallionellaceae sequences from the Silva database (1) with >1475 bp and sequence quality scores >85%. Large colored circles indicate 16S rRNA sequences from genomes in the pangenome dataset. Asterisks next to bolded names indicate 16S rRNA from isolates.

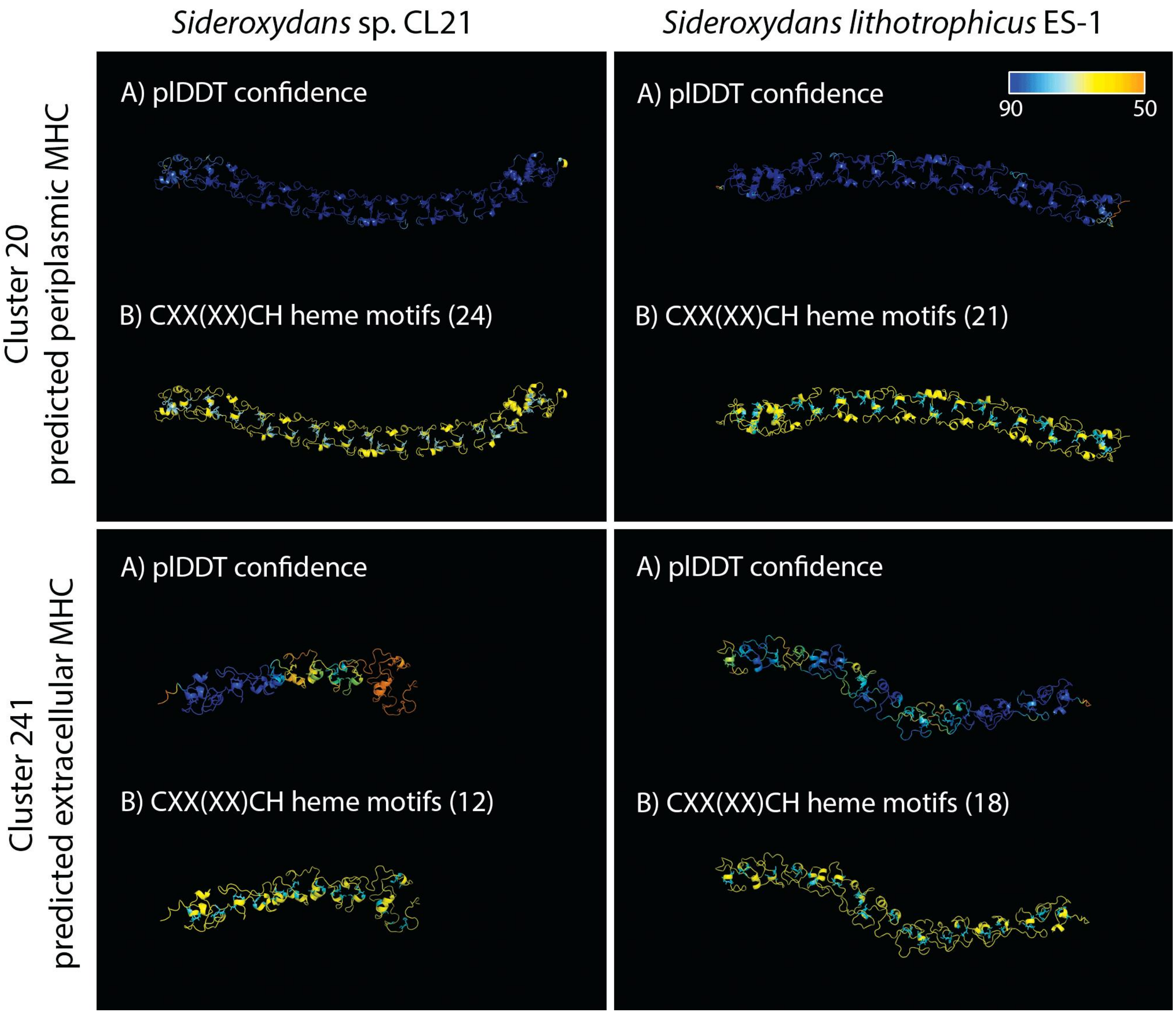


**FIGURE S2** - AlphaFold2 (2, 3) models of predicted PCC3 proteins. Proteins colored A) by plDDT confidence with dark blue representing over >90% confidence and orange representing <50% confidence, and B) with the C and CH of predicted CXX(XX)CH motifs shown in cyan to represent the position of heme-binding within the yellow protein structure.

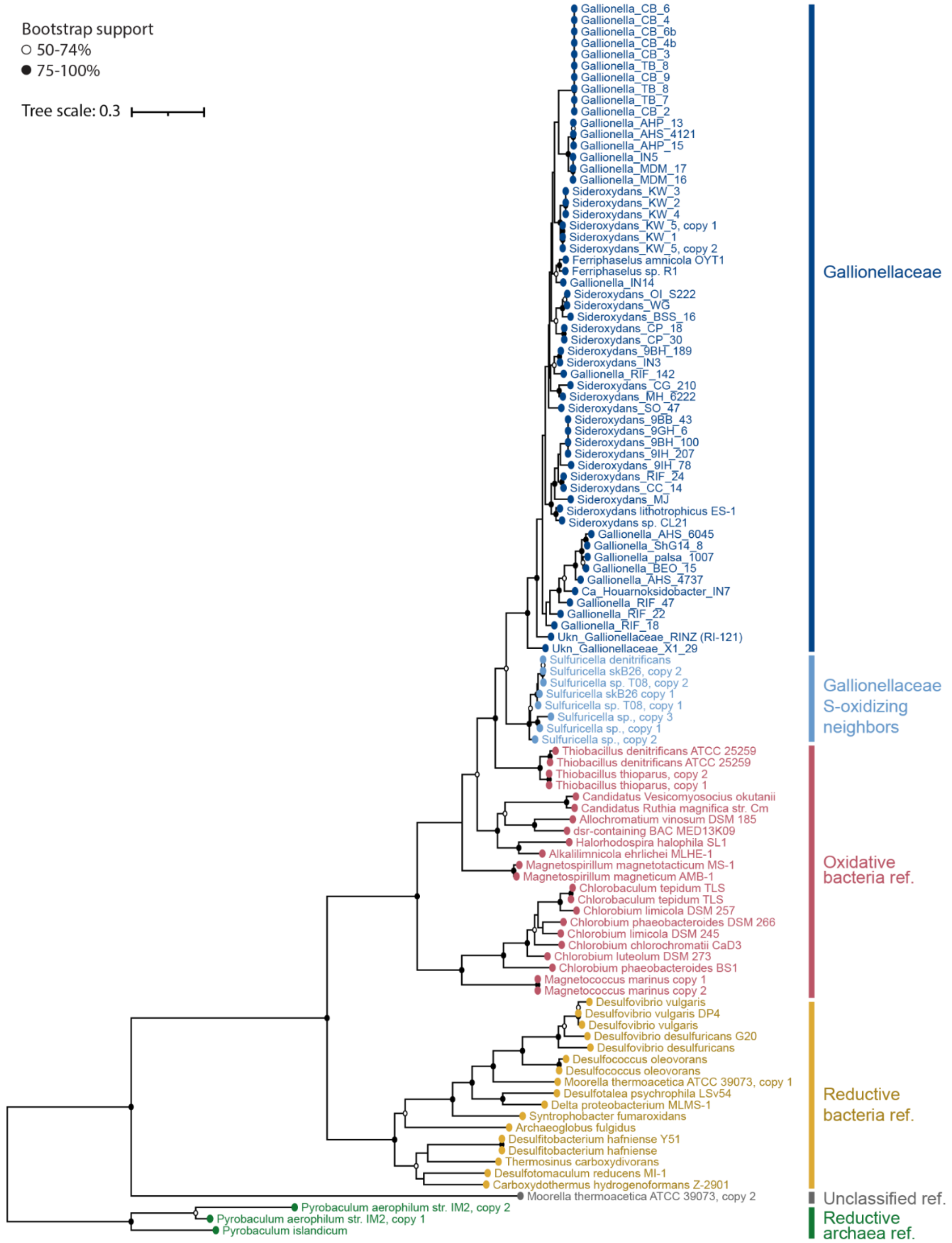


**FIGURE S3** Concatenated DsrAB tree of Gallionellaceae sequences plus oxidative and reductive DsrAB reference sequences from Loy, et al. (4) and Müller, et al. (5). The tree shows Gallionellaceae and their S-oxidizing neighbors have oxidative rDSR.

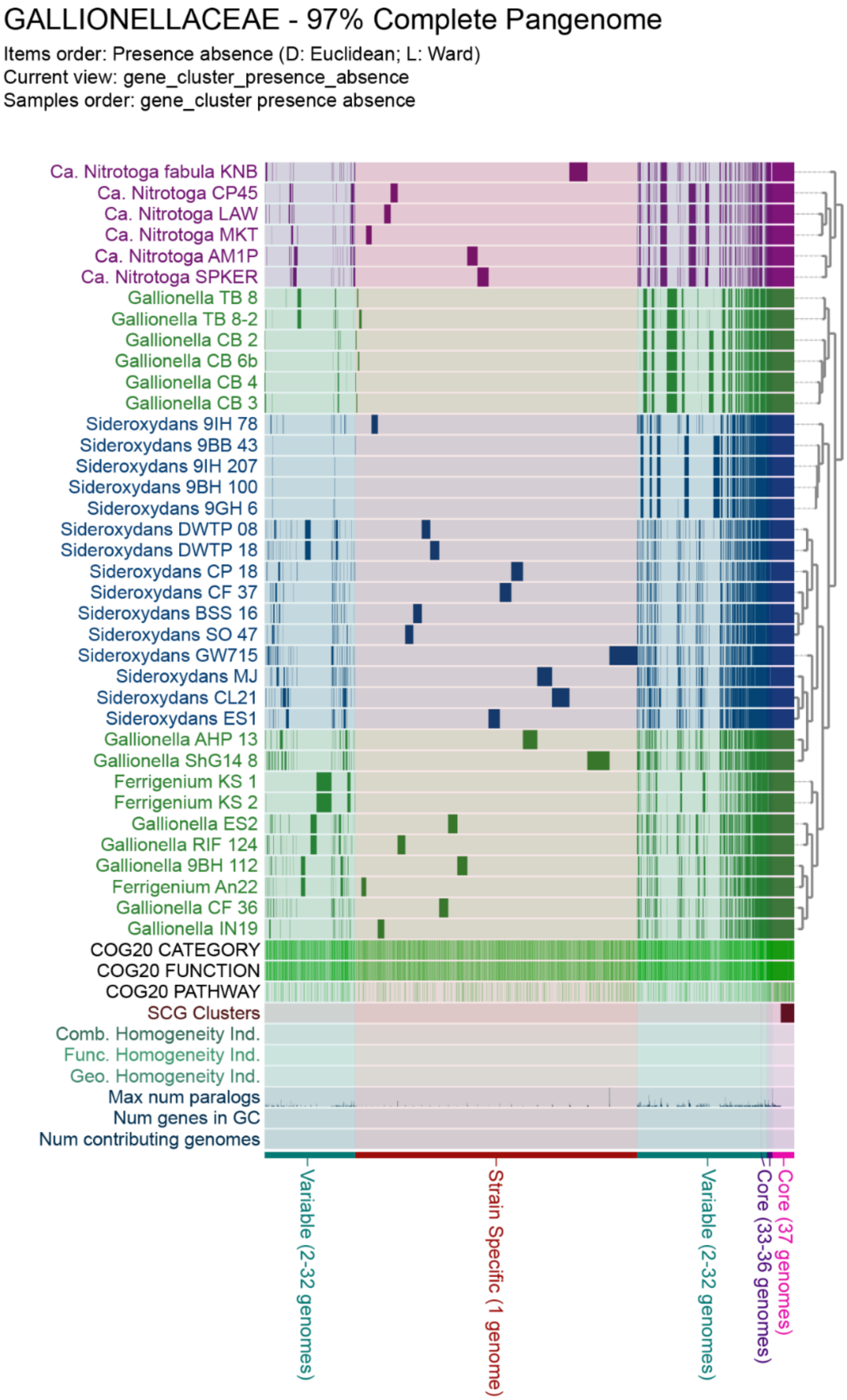


**FIGURE S4** The visual representation of the Gallionellaceae pangenome in Anvi'o (6, 7) with all genomes over 97% complete. Pangenome summary: # genomes = 37; # genes = 99,543; # gene clusters = 18,153; # gene clusters in all 37 genomes = 757 (29,404 gene calls in those clusters); # gene clusters in 33-36 genomes = 200 (7,428 gene calls in clusters); # gene clusters in 2-32 genomes (variable) = 7,531 (52,714 gene calls in those clusters); # gene clusters in 1 genome (strain specific) = 9,665 (9,997 gene calls in those clusters)

### Supplemental Tables

**TABLE S1** The table of chosen genomes with completeness, contamination, environmental and geographic metadata from databases and publications, and publication DOIs if applicable.

**TABLE S2** Table of GOLD Ecosystem Classifications.

**TABLE S3** Table of key metabolic genes.

**TABLE S4** Select Anvi'o core FeOB clusters.

**TABLE S5a and S5b** ANI and AAI matrices for the Gallionellaceae dataset.

**TABLE S6** All considered genomes with databases and accession numbers.

**TABLE S7** Table of key metabolic genes and predicted MHCs of *Sideroxyarcus emersonii*.

**TABLE S8** Naming conventions used for MAGs in this study.
